# Transcriptional and Cytotoxic Responses of Human Intestinal Organoids to Interferon Types I, II, and III

**DOI:** 10.1101/2021.12.11.472223

**Authors:** David A. Constant, Jacob A. Van Winkle, Eden VanderHoek, Simone E. Dekker, M. Anthony Sofia, Emilie Regner, Nir Modiano, V. Liana Tsikitis, Timothy J. Nice

## Abstract

The three types of interferon (IFN) have roles in antimicrobial immunity and inflammation that must be properly balanced to maintain tissue homeostasis. For example, IFNs are elevated in the context of inflammatory bowel disease and may synergize with inflammatory cytokines such as tumor necrosis factor alpha (TNFα) to promote tissue damage. Prior studies suggest that in mouse intestinal epithelial cells (IECs), type III IFNs are preferentially produced during viral infections and are less cytotoxic than type I IFN. Here, we generated human IEC organoid lines from biopsies of ileum, ascending colon, and sigmoid colon of three healthy subjects to establish the baseline responses of normal human IECs to types I, II, and III IFN. We found that all IFN types elicited responses that were qualitatively consistent across intestinal biopsy sites. However, IFN types differed in magnitude of STAT1 phosphorylation and identity of genes in their downstream transcriptional programs. Specifically, there was a core transcriptional module shared by IFN types, but types I and II IFN stimulated unique transcriptional modules beyond this core gene signature. The transcriptional modules of type I and II IFN included pro-apoptotic genes, and expression of these genes correlated with potentiation of TNFα cytotoxicity. These data define the response profiles of healthy human IEC organoids across IFN types, and suggest that cytotoxic effects mediated by TNFα in inflamed tissues may be amplified by a simultaneous high-magnitude IFN response.

## Introduction

Intestinal epithelial cells (IECs) form the primary cellular barrier to invasion by luminal microbes. Exposure to microbiota at sites of epithelial barrier damage or active invasion of the epithelium by pathogens results in antimicrobial immune responses (Constant et al., 2021). Production of cytokines such as tumor necrosis factor alpha (TNFα) and interferon (IFN) are central to an effective antimicrobial immune response. However, in the context of inflammatory bowel diseases (IBD), these cytokine responses contribute to pathology. Damage to the epithelium results in increased exposure to intestinal microbiota and chronic elevation of IFNs and TNFα, which further promote a chronic cycle of IEC barrier damage (Edelblum et al., 2006; Watson and Hughes, 2012; Blander, 2016; Okamoto and Watanabe, 2016; Friedrich et al., 2019). Additionally, IFN stimulated gene signatures have been inversely correlated with effectiveness of anti-TNFα therapy (Samie et al., 2018; Mavragani et al., 2020), suggesting that an IFN response could be a barrier to effective neutralization of TNFα in treating this disease. Therefore, it is important to understand how IEC responses to IFN balance crucial antimicrobial activities with their potential to disrupt barrier functions.

Interferons (IFNs) are a family of cytokines with pleiotropic actions in host response to pathogenic microbes, anti-tumor immunity, and autoinflammatory disease. There are three types of IFN distinguished by use of specific cellular receptors. Type I IFNs (hereafter IFN-I) signal through the IFN alpha receptor, and human IFN-I encompasses at least 13 IFN-α subtypes, IFN-β, IFN-ε, IFN-κ, and IFN-ω. Type II IFN (hereafter IFN-II) signals through the IFN gamma receptor, and is represented by a single IFN-γ gene. Type III IFNs (hereafter IFN-III) signal through the IFN lambda receptor, and human IFN-III encompasses three to four IFN-λ genes. Production of IFNs I and III is stimulated directly by sensing of pathogen-associated patterns in a wide variety of cell types, whereas IFN-II is primarily produced by activated T cells and natural killer cells. Most cell types can produce and respond to IFN-I, but IFN-III is preferentially produced by virally infected epithelial cells and has a prominent role in control of viral infection in the intestinal epithelium (Mordstein et al., 2010; Pott et al., 2011; Odendall et al., 2014; Mahlakoiv et al., 2015; Nice et al., 2015; Baldridge et al., 2017).

IFNs stimulate gene expression via activation of Janus kinases (JAKs) and subsequent phosphorylation of signal transducer and activator of transcription (STAT) factors (Villarino et al., 2017). There are multiple STATs that can be activated by IFN signaling and determine the scope of downstream gene expression (Au-Yeung et al., 2013). STAT1 is phosphorylated downstream of all three IFN types, and is the predominant STAT family member activated by IFN-II. Phosphorylated STAT1 dimerizes and transactivates expression of IFN-stimulated genes (ISGs) that have a consensus promoter motif called the interferon-gamma-activated sequence (GAS) element. Canonical signaling by IFN-I and IFN-III result in phosphorylation of STAT2 as well as STAT1, which join with IFN regulatory factor 9 (IRF9) to form a heterotrimeric transcription complex called interferon stimulated gene factor 3 (ISGF3). ISGF3 binds to interferon stimulated response elements (ISREs) in ISG promoters and can independently activate transcription or co-operate with transcriptional activators of other motifs, such as GAS elements. There are hundreds of ISGs, each with a distinct arrangement of promoter elements, including some with IFN-type-specific and cell-type-specific expression patterns (Schoggins, 2019).

Development of organoid culture systems has significantly advanced studies of primary IEC biology (Sato et al., 2009; Miyoshi and Stappenbeck, 2013). IEC organoids are derived from crypt-resident stem cells that can be isolated from intestinal biopsy and are maintained by inclusion of stem cell maintenance factors. These IEC organoid systems are particularly well suited for defining intrinsic cytokine responses “hard-wired” within the IEC lineage that operate independently from the complex intestinal milieu. We previously studied IFN responses of mouse IEC organoids and found that IFN-III responses are highly conserved between IEC organoids and IECs *in vivo* (Van Winkle et al., 2020). We also found that IFN-I elicits robust ISG expression in IEC organoids, with increased magnitude of response compared to IFN-III. IFN-I stimulated greater overall expression of shared ISGs and stimulated a greater total number of ISGs in mouse IEC organoids This expanded ISG response to IFN-I included pro-apoptotic factors, and IFN-I pretreatment resulted in potentiation of TNFα-induced IEC cytotoxicity to a greater extent than IFN-III (Van Winkle et al., 2020).

In the present study, we sought to extend our prior observations of IFN-potentiated TNFα cytotoxicity mouse IECs to human IEC organoids derived from multiple intestinal biopsy sites and subjects and to all three types of IFN. We find that combined IFN treatment consistently potentiates TNFα-induced IEC cytotoxicity in all IEC organoid lines tested. To characterize IFN type-specific responses, we performed dose-response analyses of IFN treatments and found that human IECs responded to IFN-I with increased phosphorylation of STAT1 and increased fold-change of ISGs compared to IFN-III. Additionally, we found that of the three tested IFN types, IFN-II was the most potent stimulator of STAT1 phosphorylation, and stimulated expression of ISGs shared with IFN-I as well as a unique ISG signature. Pro-apoptotic genes were stimulated by IFN-II and IFN-I to a greater degree than IFN-III, and expression of these “apoptosis ISGs” correlated with potentiation of TNFα-induced cytotoxicity. These findings emphasize that regulation of IFN response magnitude and selectivity among IFN types is crucial to balancing antimicrobial actions with pathological potential.

## Results

### Generation of IEC organoids from healthy human subjects

We generated IEC organoids from intestinal biopsies of healthy subjects undergoing colonoscopies at OHSU. Biopsies were collected from terminal ileum, ascending colon, or sigmoid colon, and organoids were generated from these tissues according to established procedures (**Fig. 1**) (Miyoshi and Stappenbeck, 2013; VanDussen et al., 2015). We selected organoid lines derived from three healthy subjects to characterize the normal cytokine responses of IECs from these tissue sites and assess the degree of variance between independently generated healthy IEC organoid lines. These subjects represented a range of participant ages, including a 20-year-old male (subject 005), a 52-year-old female (subject 007), and a 71-year-old male (subject 008).

**Figure 1.**
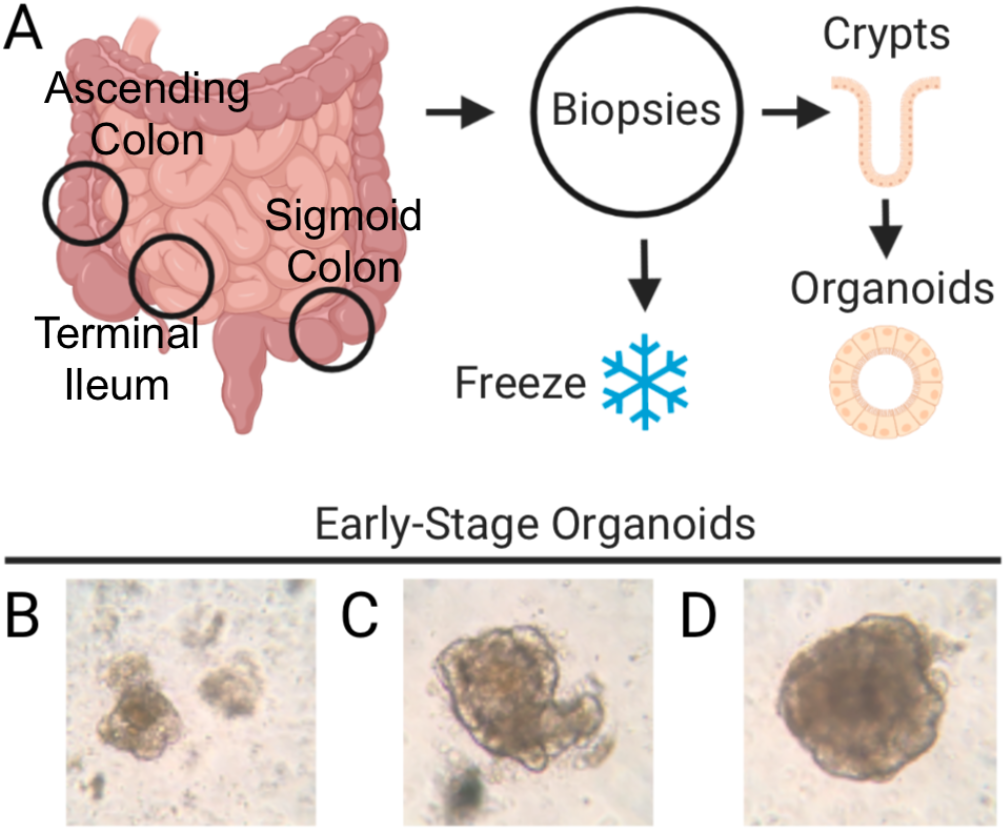
Sample collection and organoid generation. **A)** Diagram of biopsy sites and processing. Biopsies were taken from terminal ileum, ascending colon, and sigmoid colon. Isolated crypt preparations were plated in Matrigel and cultured with stem cell maintenance factors to generate IEC organoids. **B-D)** Representative images of newly formed (P0) organoids derived from the terminal ileum **(B)**, ascending colon **(C)**, and sigmoid colon **(D)**. Micrographs were taken at 10X magnification on a phase-contrast dissecting microscope.

### Combined IFN treatment potentiates TNFα cytotoxicity in human IEC organoids

TNFα can stimulate growth or death of IECs depending on the cellular context and expression of pro-apoptotic factors (Leppkes et al., 2014; Kalliolias and Ivashkiv, 2016). Based on our previous findings in mouse IECs (Van Winkle et al., 2020), we expected that an IFN response in human IEC organoids would potentiate TNFα cytotoxicity. To test this hypothesis, we treated IEC organoids with pooled IFN types I-III to stimulate a concerted IFN pathway response, and subsequently exposed the IEC cultures to TNFα. For comparison, we included conditions where organoids were left untreated, IFN-treated alone, or TNFα-treated alone. Visualization of apoptotic cells using cleaved caspase 3 indicated that TNFα-treatment alone resulted in minimal apoptosis, and IFN-treatment alone stimulated a modest amount of apoptosis (**Fig. 2A-B**). However, combined TNFα and IFN treatment resulted in cooperative induction of apoptosis that was significantly increased from either treatment alone (**Fig. 2A-B**).

**Figure 2.**
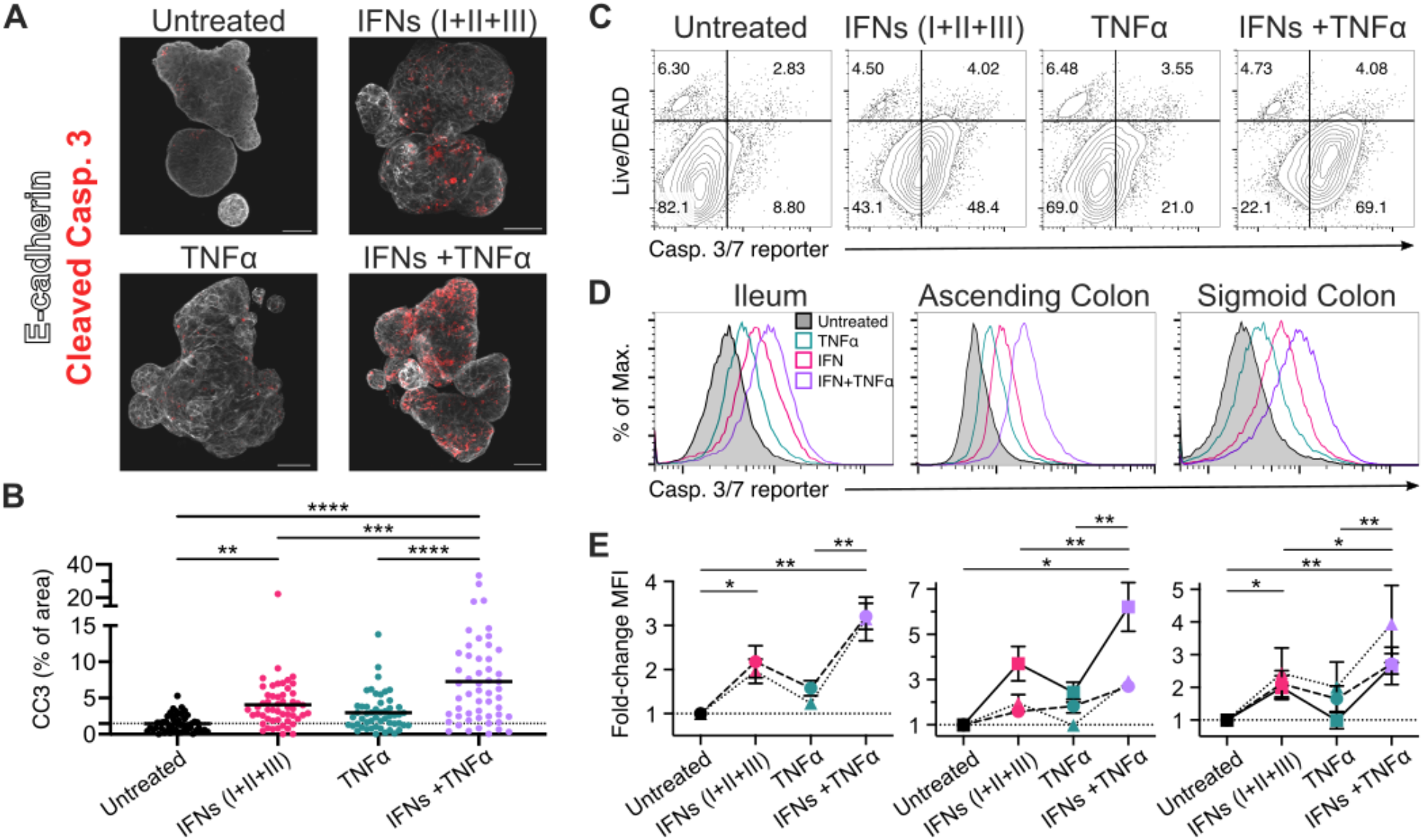
Combined IFN treatment potentiates TNFα cytotoxicity in human IEC organoids. IEC organoids were treated for 4 hours with PBS or pooled IFN types I, II, and III (1,000 U/mL each), followed by treatment with PBS or TNFα (100 ng/mL) for 20 hours. **A)** Representative immunofluorescent micrographs of IEC organoids after the indicated treatments, showing cleaved caspase 3 (red) and E-cadherin (white), scale bar = 50µm. **B)** Summary data of two independent experiments using organoids from subjects 005 and 008. Data shows percent area positive for cleaved caspase 3 (CC3) relative to the E-cadherin positive area per field of view. Total fields of view: untreated, N=31; IFNs, N=32; TNFα, N=30; IFNs +TNFα, N=32; line indicates mean. **C)** Representative flow cytometry plots of organoids subjected to the indicated treatments and stained with a caspase 3/7 reporter and “live/DEAD” cell permeability dye. **D)** Representative histograms showing the median fluorescence intensity (MFI) of caspase 3/7 activity reporter among intact IECs (negative for live/DEAD dye) in response to the indicated treatments. **E)** Summary data of fold-change in caspase 3/7 reporter fluorescence relative to matched untreated control organoids. Subject 005 = dashed line; subject 007 = solid line; subject 008 = dotted line. Data in **(E)** represents mean of three independent experiments. Statistical significance determined by one-way ANOVA **(B)** or two-way ANOVA **(E)** with Tukey’s multiple comparison test. * = p<0.05; ** = p <0.01; *** = p<0.001; **** = p < 0.0001.

To compare this cooperative cytotoxic response in a more sensitive and high-throughput manner across healthy subjects and tissue sites, we used a caspase 3/7 activity reporter to analyze apoptosis in dissociated IEC organoids by flow cytometry. By inclusion of a live/dead discrimination dye, we found that the majority of cells retained membrane integrity at this timepoint, with a minor proportion of cells lysed during organoid dissociation (∼5-10% Live/DEAD-positive in Untreated, **Fig. 2C**). For all experimental groups, we observed minimal increase in lysed cells and an increase in caspase 3/7 activity within the intact cell population (**Fig. 2C**). Therefore, we quantified apoptosis stimulation as the fold-change in fluorescence intensity of caspase 3/7 reporter within intact cells relative to untreated controls (**Fig. 2D-E**). There was a statistically significant increase in caspase 3/7 activity resulting from combined IFN and TNFα treatment across all sites and subjects tested (**Fig. 2E**). Treatment with pooled IFNs alone also resulted in a statistically significant increase in caspase 3/7 activity, but to a lesser degree than the combination treatment, and TNFα alone caused a modest increase in caspase 3/7 activity that was not statistically significant (**Fig. 2E**). Together, these data indicated cooperative induction of apoptosis between IFN and TNFα occurs in human IECs, with phenotypic consistency across organoids generated from multiple subjects and tissue sites.

### Differential stimulation of STAT1 phosphorylation by IFN types in IEC organoids

To characterize the responses of IEC organoids to individual IFN types, we used a high-throughput assay to detect STAT1 phosphorylation (pSTAT1) by time-resolved fluorescence resonance energy transfer (TR-FRET). We treated cells with 100-fold dilutions of purified IFN types I, I, or III to compare the extent of pSTAT1 elicited by each IFN type (**Fig. 3A-B**). We found that IFN-II induced the strongest maximal pSTAT1 response and TR-FRET signal was detectable from as little as 1 U/mL, but IFN-III stimulated marginal pSTAT1 at the highest 10,000 U/mL dose. IFN-I induced an intermediate pSTAT1 response, and was detectable beginning at the 100 U/mL dose. This hierarchy of pSTAT1 stimulation between IFN types was consistent across IEC organoids from different subjects and tissue sites, but the most uniform responses between subjects were observed in organoids from sigmoid colon (**Fig. 3B**). Overall, these data suggest that a similar pattern of responsiveness to IFN types is present across IEC organoids from different subjects and tissue sites, with IFN-II consistently stimulating the most potent phosphorylation of STAT1.

**Figure 3.**
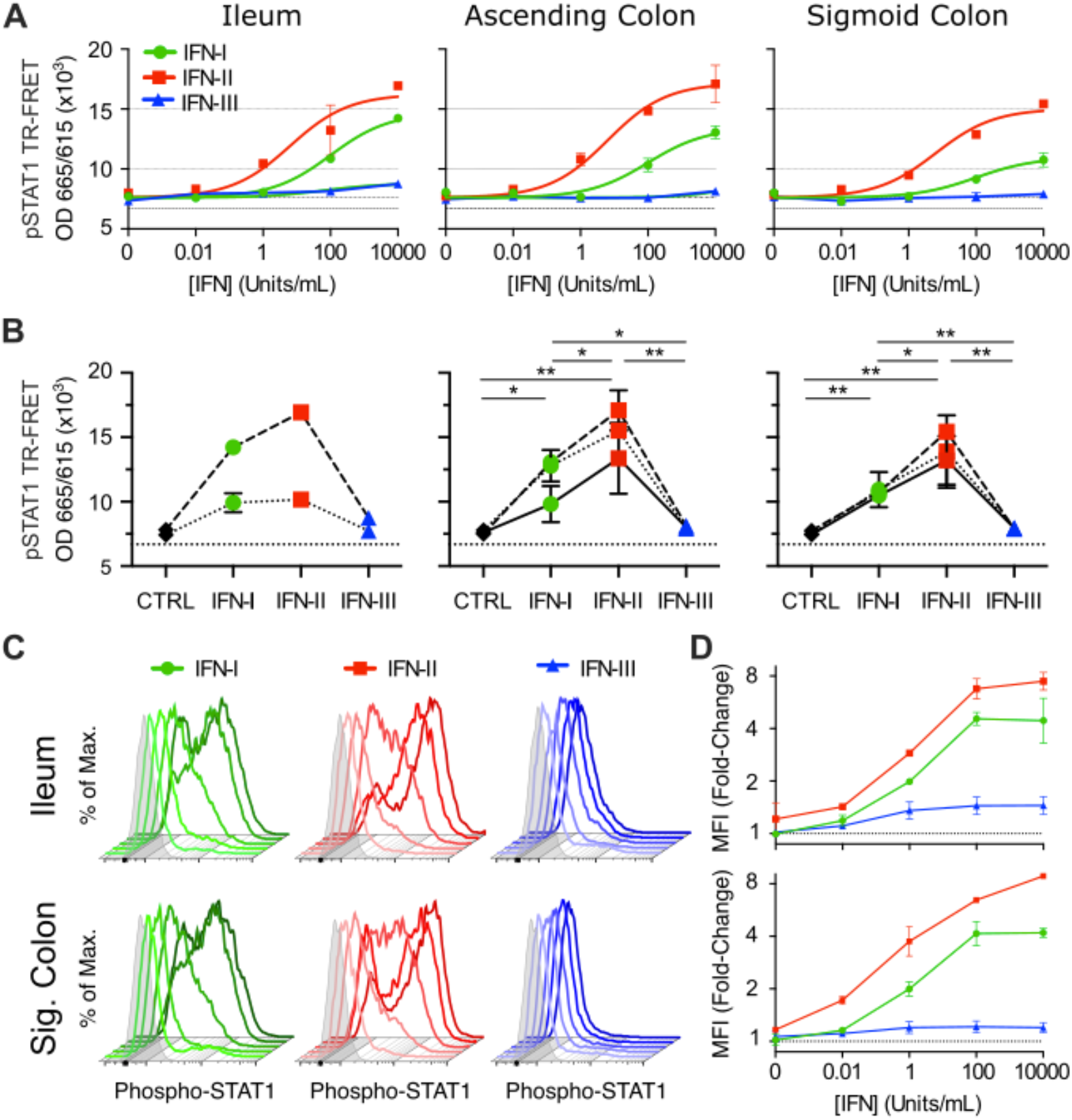
Dose-response of STAT1 phosphorylation in IEC organoids. IEC organoids were treated with titrations of 1-10,000 U/mL of type I, type II, or type III IFN for one hour and pSTAT1 (Y701) levels were measured by TR-FRET (**A-B**) or flow cytometry (**C-D**). **A)** Representative dose-response curves are shown for organoids from ileum, ascending colon, and sigmoid colon of subject 005. **B))** Comparison of maximal pSTAT1 detected by TR-FRET for indicated IFN types, subjects, and tissue sites. Dashed line indicates subject 005, solid line subject 007, and dotted line subject 008. **C)** Representative histograms from flow cytometry detection of pSTAT1 levels for ileum and sigmoid colon of subject 005. Increasing IFN concentrations are indicated by increasing darkness of histogram lines. **D)** Data from ileum and sigmoid colon for all conditions are summarized. Data in A, B, and D are means with SEM from two biological replicates. Significance determined by two-way ANOVA with Tukey’s multiple comparison test. * = p<0.05; ** = p <0.01; *** = p<0.001.

The prior TR-FRET data represented an average response within the organoid cultures, but it remained possible that there were heterogeneous responses between individual IECs within a culture, as had been previously reported in an IEC cell line (Bhushal et al., 2017). Therefore, we extended our analysis using a more sensitive and quantitative flow cytometry measurement of pSTAT1. We selected a single subject for flow cytometry analysis of ileum and sigmoid colon organoids treated with titrations of IFN types I-III as above. In all cases, we found that the pSTAT1 response was consistently observed in >80% of IECs relative to background pSTAT1 staining (**Fig. 3C**), indicating that the majority of IECs within organoid cultures can respond to IFN stimulation. However, at intermediate doses of IFN-II or IFN-I, there was heterogeneity in pSTAT1 intensity among responding IECs that could reflect stochastic effects or indicate a subset of IECs with differential responsiveness. Similar to TR-FRET results, we found that the highest magnitude of pSTAT1 was stimulated by IFN-II, with intermediate stimulation by IFN-I and the lowest magnitude stimulated by IFN-III (**Fig. 3D**). IFN-III stimulated only a modest 20-50% increase in pSTAT1 fluorescence above background levels, but was detectable beginning from the one U/mL dose that was not apparent by TR-FRET, reflecting an increased sensitivity to pSTAT1 detection by flow cytometry. These results suggest the majority of IECs within the cultured organoids are similarly capable of responding to each IFN type. Additionally, the presence of a pSTAT1 response to IFN-III is confirmed, but is clearly of much lower magnitude relative to IFN-I and IFN-II.

### Differential ISG response between IFN types in IEC organoids

We extended our analysis of IEC organoid responses to the downstream transcriptional activation of representative ISGs (*IRF1, CXCL10*, and *ISG15*). IRF1 is a transcription factor that can provide additional positive-feedback stimulation of ISGs, CXCL10 is a secreted chemokine, and ISG15 is a known antiviral effector. These three ISG promoter regions contain GAS elements (*IRF1*), ISRE motifs (*ISG15*), or both (*CXCL10*) (**Fig. 4A**), and their expression was therefore predicted to differentially depend on STAT1 homodimers and ISGF3.

**Figure 4.**
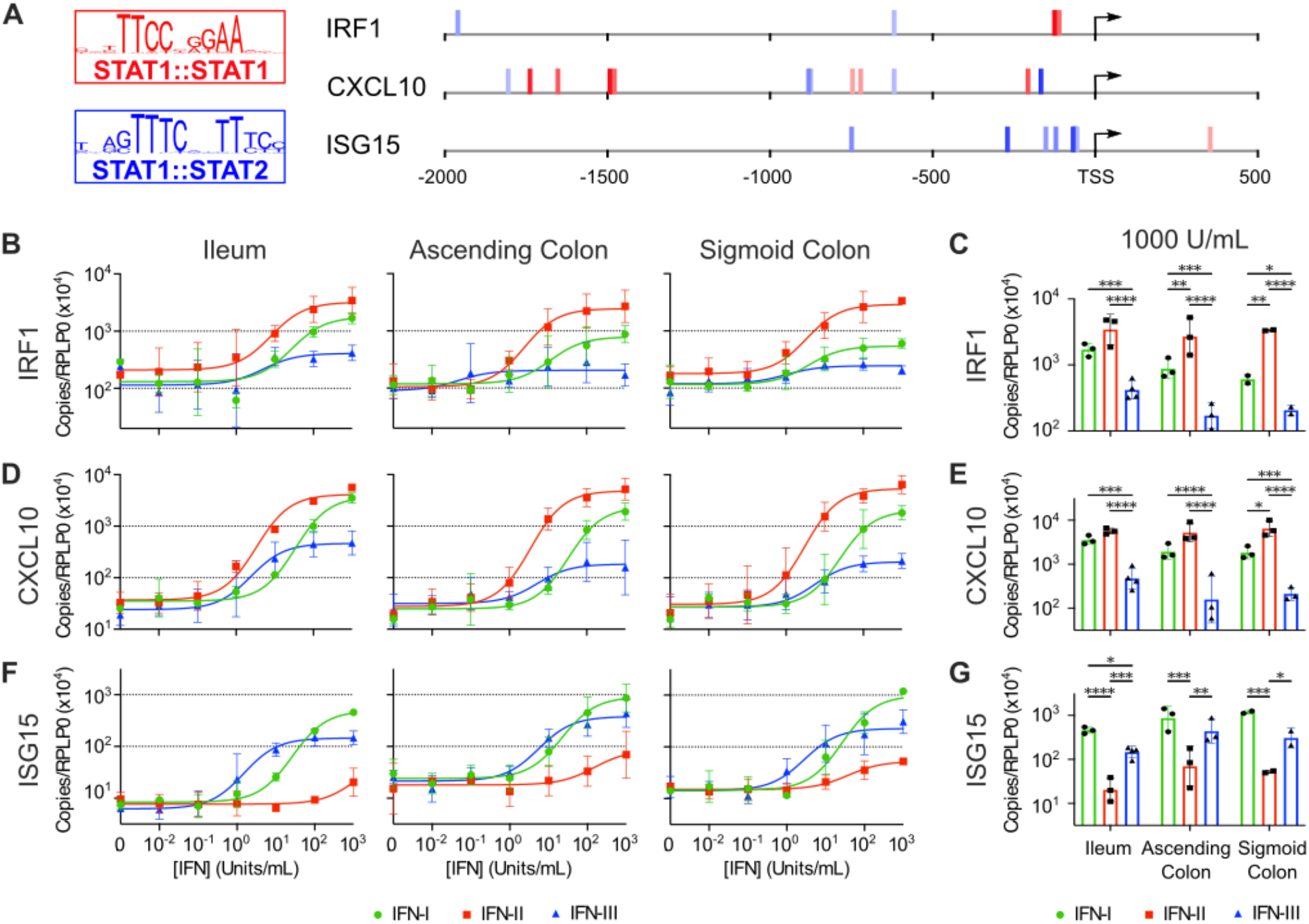
Dose-response of ISG transcription in IEC organoids. **A)** GAS elements (red) and ISREs (blue) in promoter regions of *CXCL10, IRF1*, and *ISG15*. Motifs with score greater than five are shown, with darker shading of boxes indicating higher-scoring motifs. **B-G)** Organoids from ileum, ascending colon, and sigmoid colon of subject 005 were treated with titrations of IFN-I, IFN-II, or IFN-III as shown, and expression of *CXCL10* (**B-C**), *IRF1* (**D-E**), and *ISG15* (**F-G**) was measured by qPCR. Data shown are normalized to expression of *RPLP0* and are from 3-4 independent experiments. Significance determined by two-way ANOVA with Tukey’s multiple comparison test. * = p<0.05; ** = p <0.01; *** = p<0.001; **** = p<0.0001.

We treated organoids from ileum, ascending colon, and sigmoid colon of subject 005 with titrations of IFN-I, IFN-II, or IFN-III, and observed that *CXCL10* and *IRF1* were stimulated to the greatest extent by IFN-II treatment, with marginal stimulation of these ISGs by IFN-III and intermediate stimulation by IFN-I (**Fig. 4B-E**). This hierarchy of gene transactivation was correlated to the pSTAT1 response elicited by these IFN types and consistent with the presence of GAS elements in these gene promoters. In contrast, *ISG15* was most robustly stimulated by IFN-III and IFN-I, with modest stimulation by IFN-II (**Fig. 4F-G**). Together, these data showed stimulation of representative ISGs by all IFN types, with magnitude of stimulation differing among IFN types and correlating with presence of promoter motifs.

To comprehensively identify differential ISG stimulation across IFN types in human IECs, we performed RNA sequencing using the IFN dose that elicited maximum expression of *IRF1, CXCL10*, and *ISG15* (1000 U/mL, **Fig. 4**). We selected three organoid lines from two healthy subjects (subjects 005 and 007) to facilitate identification of normal ISG signatures that were consistent between IEC lines generated from independent healthy human biopsies. Principal component analysis of this experiment revealed the primary separation of samples (PC1, 75% variance) was related to the subject and site from which the IEC organoid line was generated (**Fig. 5A**). This is consistent with prior studies showing that ileum and colon IECs have distinguishing epigenetic features that are retained in organoid culture (Kraiczy et al., 2019), and emphasizes the importance of including multiple organoid lines in these analyses. Even so, IFN treatment groups for each IEC organoid line were indicated by a secondary component (PC2, 9% variance, **Fig. 5A**). Furthermore, read counts from the RNAseq dataset for *IRF1, CXCL10*, and *ISG15* were increased in IFN treated cells, and the IFN-type-specific hierarchies were identical to qPCR (**Fig. 5B)**.

**Figure 5.**
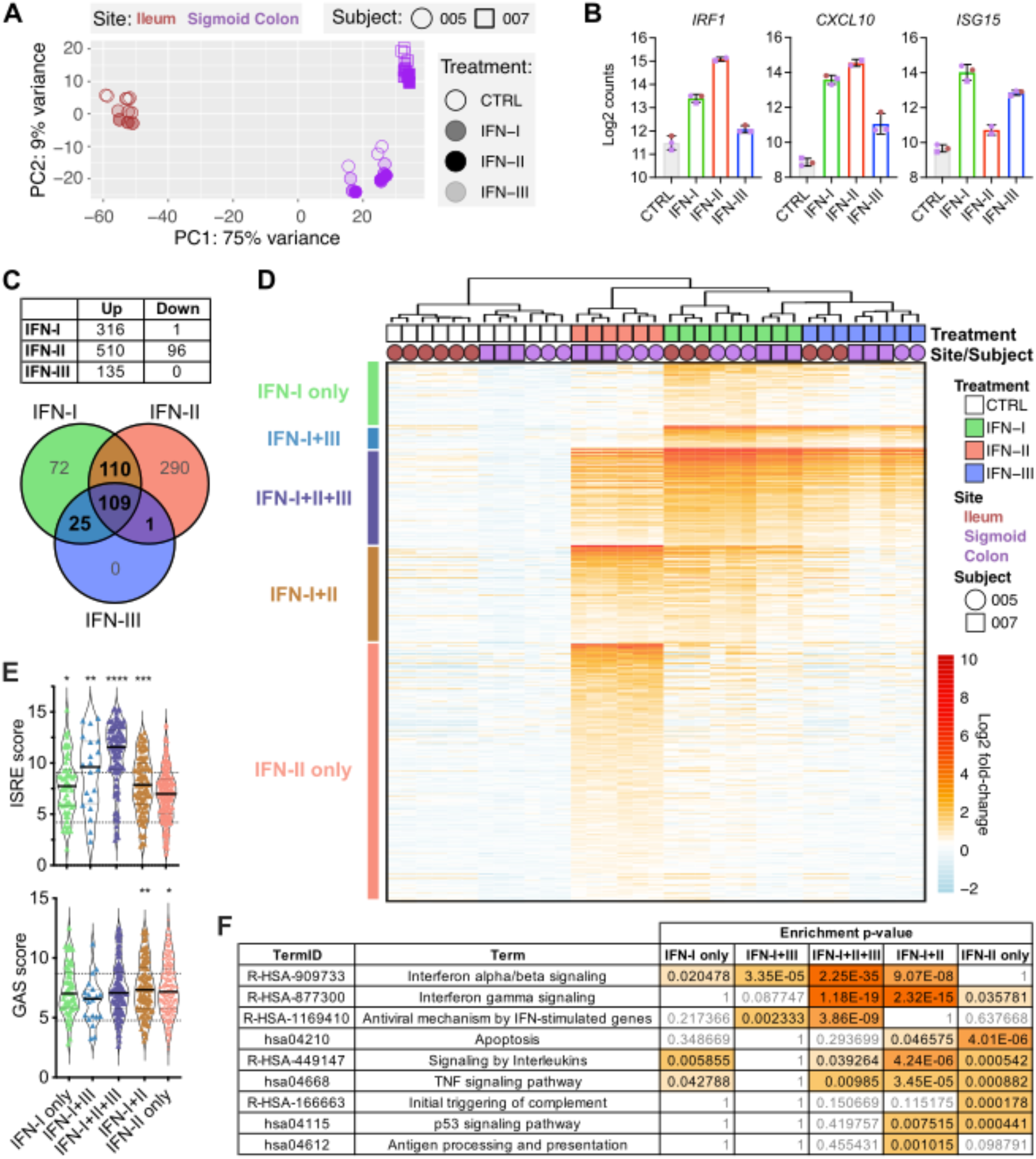
Definition of ISG modules in IEC organoids by RNAseq. Organoids from ileum of subject 005, sigmoid colon of subject 005, and sigmoid colon of subject 007 were treated with 1000 U/mL of IFN-I, IFN-II, or IFN-III for four hours and RNA was isolated for RNA sequencing. **A)** Principal component analysis of all samples. **B)** Log2 counts for the indicated ISGs. **C)** Table shows number of genes upregulated and downregulated by indicated IFN treatments relative to control, and Venn diagram shows overlap of upregulated genes across IFN types. **D)** Heatmap of log2 fold-change in counts for each ISG relative to mean value of control samples. Samples are clustered according to the dendrogram and genes are arranged by ISG modules, as indicated by colored bars at left of heatmap. **E)** Maximum ISRE and GAS motif scores within promoter regions (−2000 to +500 relative to TSS) of each gene within the indicated ISG modules. **F)** Selected pathways that are differentially enriched among ISG modules and associated p-values.

All IFN treatments stimulated expression of hundreds of genes relative to control treatments, with relatively few down-regulated genes. We defined ISGs in this dataset as genes with greater than 1.5-fold increase relative to control treatment and an adjusted p-value of < 0.05. Using these criteria, IFN I, II, and III treatments were associated with 316, 510, and 135 ISGs, respectively (**Fig. 5C**). There was substantial overlap between IFN types, with a “core” signature of 245 ISGs shared by at least two IFN types and 109 ISGs shared across all three IFN types. However, there were also genes unique to individual IFN types, with 72 ISGs significantly stimulated by IFN-I only and 290 ISGs significantly stimulated by IFN-II only (**Fig. 5C-D**). IFN-III did not stimulate any unique ISGs, with 134/135 genes shared with IFN-I and 1/135 genes shared with IFN-II. Together, these comparisons defined modules of shared and unique ISGs across each IFN type and three healthy IEC organoid lines.

Analysis of canonical promoter elements present within shared and unique ISG modules confirmed that ISRE elements were significantly enriched in promoters of IFN-I-stimulated genes, and GAS elements were significantly enriched in promoters of IFN-II-stimulated genes (**Fig. 5E**). Furthermore, ISRE scores of genes in the “IFN-I only” ISG module were lower than ISRE scores of genes shared between IFN-I and IFN-III (“IFN-I+III” and “IFN-I+II+III” modules, **Fig. 5E**). These data indicate that IFN-I can transactivate promoters that lack high-scoring canonical ISRE elements, and suggest that IFN-III is more selective in transactivation of promoters containing high-scoring canonical ISRE elements.

To gather physiological information about ISG modules, we tested for enrichment of pathway genesets curated by KEGG and Reactome databases, and identified pathways of interest that were differentially enriched across modules (**Fig. 5F**). The “Interferon alpha/beta signaling” and “Interferon gamma signaling” Reactome pathways were represented to varying extents across modules, but the greatest enrichment of both pathways was in the core “IFN-I+II+III” module, followed by the shared “IFN-I+II” module. The “Antiviral mechanism by IFN-stimulated genes” Reactome pathway was significantly enriched in modules shared by IFN-I and IFN-III (“IFN-I+III” and “IFN-I+II+III”), but not in the remaining modules (**Fig. 5F**). In contrast, the “Apoptosis” and “p53 signaling” KEGG pathways were significantly enriched in “IFN-II only” and “IFN-I+II” modules but not other modules. Few curated pathways were associated with genes in the “IFN-I only” module, but among these were the “Signaling by Interleukins” Reactome pathway and the “TNF signaling pathway” KEGG pathway, which were also highly enriched in the “IFN-I+II” ISG module (**Fig. 5F**). This pathway information points to differing biological functions of the IFN types, and is consistent with a targeted role for IFN-III in stimulating antiviral genes and broader roles of IFNs I and II associated with more expansive modules of ISG expression.

### Differential potentiation of TNFα cytotoxicity by IFN types

To more specifically hone in on ISGs related to apoptosis, we identified genes in the KEGG apoptosis pathway that were present among the ISG modules identified by RNAseq. These “apoptosis ISGs” were predominantly pro-apoptotic genes (including *BAK1, CASP3, CASP8, CASP10, FAS, TNFSF10A*, and *TNFRSF10A*), and were stimulated by IFN-II or IFN-I treatments (**Fig. 6A**). The preferential stimulation of apoptosis ISGs by IFN-II and IFN-I is consistent with lower p-values of KEGG apoptosis pathway enrichment within “IFN-I+II” and IFN-II only” ISG modules.

**Figure 6.**
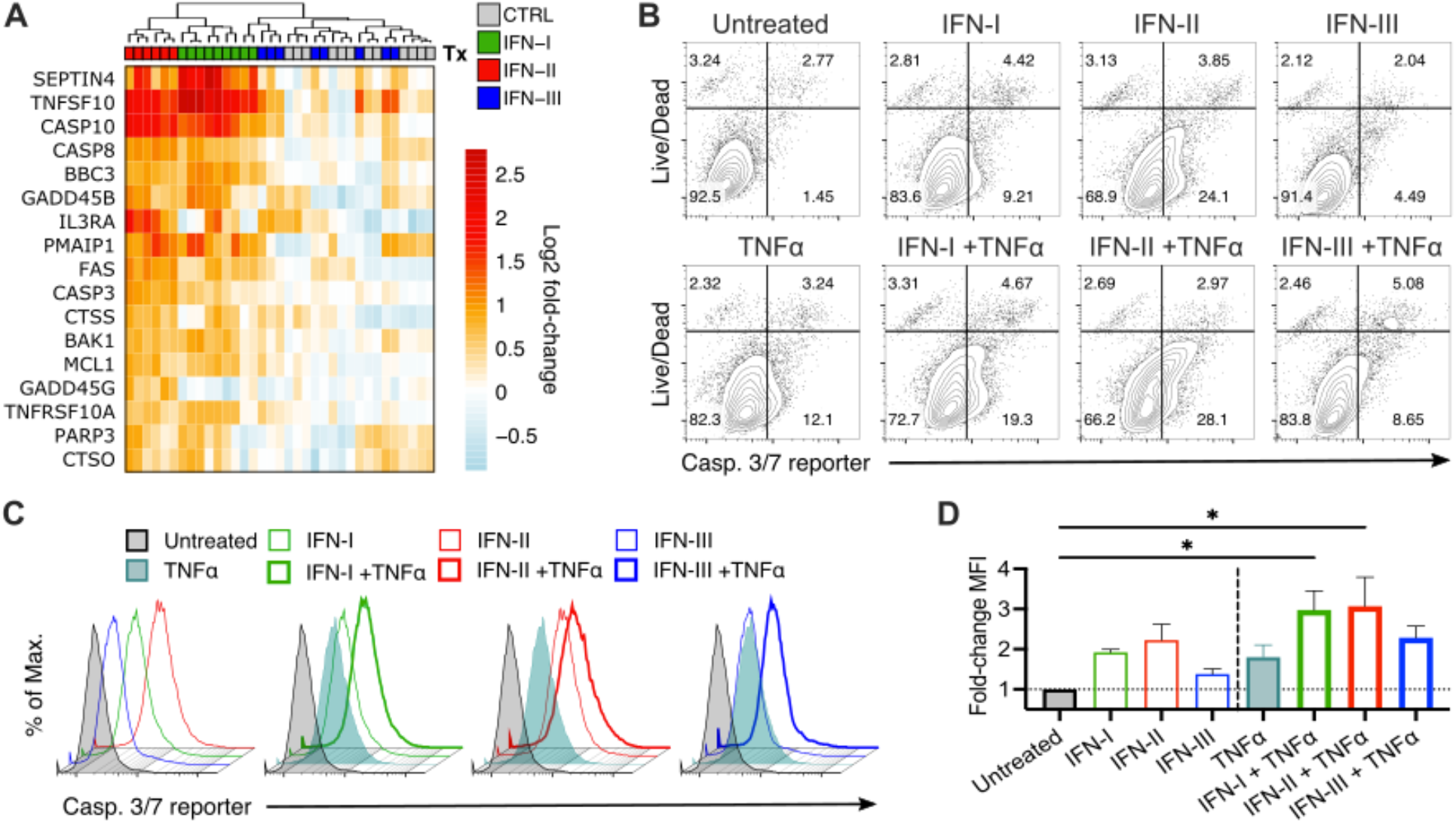
Differential potentiation of TNFα cytotoxicity by IFN types. **A)** Fold-change for genes in the KEGG Apoptosis pathway (hsa04210) that are present within ISG modules from RNAseq data in figure 5. **B-D)** IEC organoids were treated for 4 hours with PBS, IFN-I, IFN-II, or IFN-III, followed by treatment with PBS or TNFα for 20 hours. **B)** Representative flow cytometry plots of organoids subjected to the indicated treatments and stained with a caspase 3/7 reporter and “live/DEAD” cell permeability dye. **C)** Representative histograms showing the median fluorescence intensity (MFI) of caspase 3/7 activity reporter among intact IECs (negative for live/DEAD dye) in response to the indicated treatments. **D)** Mean fold-change in caspase 3/7 reporter fluorescence relative to untreated organoids from four independent experiments (subjects 005 and 008) with error bars indicating SEM. Statistical significance determined by one-way ANOVA with Tukey’s multiple comparison test. * = p<0.05.

To test whether increased expression of apoptosis ISGs between IFN types resulted in functional differences in activation of apoptosis in IEC organoids, we compared caspase 3/7 activity following individual IFN treatments. We found that IFN-II and IFN-I stimulated a greater increase in caspase 3/7 activity than IFN-III **(Fig. 6B-D**). Additionally, IFN-II or IFN-I treatments were sufficient to increase the apoptotic potential of TNFα, but IFN-III treatment resulted in minimal increase in caspase 3/7 activity compared to TNFα alone **(Fig. 6B-D**). Together, these data indicate that IFN-I and IFN-II can cooperate with TNFα to induce apoptosis of healthy IEC organoids. This IFN type-specific effect correlates with our prior observations of increased STAT1-phosphorylation and transactivation of ISG modules beyond ISGs with canonical ISRE elements. This cooperative response between IFN responses and TNFα may contribute to loss of barrier integrity during inflammatory disease, emphasizing the importance of homeostatic regulation of IFN responses in healthy IECs.

## Discussion

Here we have shown that apoptosis of human IEC organoids in response to TNFα is potentiated by a simultaneous IFN response, and that this response is consistent among IEC lines independently generated from distinct biopsy sites and healthy donors. We additionally show that type I and II IFN are similarly able to cooperate with TNFα, but IFN-III stimulation is significantly less pro-apoptotic in healthy IECs. Our findings are consistent with prior characterizations of synergy between IFN-II and TNFα in disruption of barrier integrity or viability of IEC cell lines and organoids (Fish et al., 1999; Wang et al., 2006; Woznicki et al., 2020), and indicate that this property can be shared between types I and II IFN.

We have confirmed that primary human IECs are responsive to all three IFN types, and find that a core signature of antiviral ISGs with high-scoring ISRE motifs is stimulated by all IFN types. IFN-III is particularly selective for stimulation of these core genes, which are enriched for antiviral effectors, with no unique ISGs that are not stimulated by other types. However, IFN-I and IFN-II stimulate unique modules of ISGs, including many ISGs with low-scoring ISRE motifs. Pathway analysis indicates that IFN-I and IFN-II stimulate pathways beyond the core antiviral effector genes, including p53 signaling, TNF signaling, and antigen processing. The expanded ISG signature elicited by types I and II IFN also include pro-apoptotic genes such as *CASP8, CASP3*, and *BAK1* that may underlie the increased apoptotic cell death stimulated by TNFα. It is likely that these intracellular pro-apoptotic factors operate in concert with additional extrinsic death pathway factors such as FAS, TNF-related apoptosis-inducing ligand (TRAIL, *TNFSF10A* gene) and the TRAIL receptor (*TNFRSF10A* gene) to sensitize IEC organoid death.

Our prior studies of mouse IEC organoids showed that most ‘apoptosis ISGs’ have low-scoring canonical promoter motifs, but their stimulation by IFN-I remains dependent on STAT1 (Van Winkle et al., 2020). This led to the hypothesis that the increased number of ISGs following IFN-I stimulation relative to IFN-III is directly related to a more potent activation of STATs and increased transactivation of lower-scoring promoter motifs. Our new findings here are consistent with this model; we show that differing magnitude of STAT1 phosphorylation elicited by IFN types correlates with an expanded number of ISGs and potentiation of apoptosis.

Several recent studies of IBD reported an inverse correlation of anti-TNFα efficacy with presence of an ISG signature (Samie et al., 2018; Mavragani et al., 2020). These reports suggest that IFN responses decrease the efficacy of TNFα blockade, and may relate to the role of IFNs in amplifying epithelial apoptosis. Although IFN-III shows minimal cooperativity with TNFα in our studies of healthy IEC organoids, there is some evidence that this IFN type could develop a pathological role in IBD (Wallace et al., 2021). Our model anticipates that an IBD-associated increase in the magnitude of epithelial responsiveness to IFN (e.g. through increased expression of STAT1) may contribute to pathological responses not present in healthy tissues. It will be of interest to expand these studies to IEC organoids from IBD patients to determine whether stable differences in IFN response are associated with disease, and how treatment of IBD patients with therapeutic JAK-STAT inhibitors alters these responses. These studies lay the groundwork for future efforts to understand how IFN responsiveness is regulated in health and disease.

## Methods

### Biopsy collection, organoid generation, and organoid maintenance

Biopsies were collected from consenting adult patients at OHSU hospital from the terminal ileum, ascending colon, and sigmoid colon. Biopsies were placed in wash medium (Advanced DMEM/F12 (Gibco, #12634010) plus 10% fetal bovine serum (FBS), 1 U/mL penicillin/streptomycin with glutamate, and 1 mM HEPES) on ice and transported to the research lab. Biopsies were processed the same day to generate organoids. This study was approved by the OHSU Institutional Review Board (protocol #00020848).

Organoid generation and maintenance is performed as previously described (VanDussen et al., 2015). Briefly, biopsies are minced in 2mg/mL collagenase type I solution (Gibco, #17018029) and digested for 20 minutes at 37 °C with trituration. Digested material is then filtered (70 µm) into BSA-coated 50 mL conical tubes, diluted with 9 mL wash media and centrifuged at 500 RCF for 5 minutes. Crypts are resuspended in 1 mL wash media and centrifuged at 500 RCF. Purified crypts are then resuspended in Matrigel (Corning, # 354234) and plated in 15 µL droplets in 24-well plates and maintained in 0.5 mL conditioned media (CM) from L-WRN cells (ATCC cat#CRL-3276) which contains Wnt3a, R-spondin3, and Noggin. Inhibitors of ROCK (Y-27632, SelleckChem) and TGF-β (SB-431542, SelleckChem) are added to culture media to promote cell survival. Media is replaced every two days. Organoids are passaged when they become large and dense, approximately every 4-8 days. For passage, wells are rinsed with phosphate-buffered saline plus 0.5 mM EDTA and trypsinized for 2.5-10 minutes, diluted with washing media, and centrifuged at 500 RCF for 5 minutes. Cells are resuspended in Matrigel and plated as above at 30,000-100,000 cells per well.

### Cytokines

Recombinant cytokines were purchased from indicated suppliers: IFN-β1a (Sigma, #IF014); IFN-γ (Sigma, #I17001); IFN-λ2 (IL28b; Sigma #I1288); TNFα (AbCam, #ab259410).

### Quantitative RT-PCR

RNA was isolated using the ZR *Quick*-Viral RNA or ZR *Quick*-Viral RNA 96 kit (Zymoresearch). DNA contamination was removed using the DNAfree kit (Life Technologies). cDNA was generated with the ImPromII reverse transcriptase (Promega). Quantitative PCR was performed using PerfeCTa qPCR FasMix II (QuantaBio) and the following pre-designed primer and probe assays from Integrated DNA Technologies (IDT): *CXCL10* – assay number Hs.PT.58.3790956.g; *ISG15 –* assay number Hs.PT.58.39185901.g; *IRF1 –* assay number Hs.PT.58.26847423; *RPLP0* – assay number Hs.PT.39a.22214824. Absolute copy number was determined by comparing Ct values to a standard curve generated using DNA of known copy number encoding the target sequences. Samples are graphed as absolute copy number of the indicated target divided by the absolute copy number of the housekeeping gene, *RPLP0*.

### Promoter motif analysis

Motif position weighted matrix files were obtained from the JASPAR database for GAS (MA0137.2) and ISRE (MA0517.1) elements (Hartman et al., 2005; Robertson et al., 2007). The findMotifs.pl function from the Hypergeometric Optimization of Motif EnRichment (HOMER) software package (Heinz et al., 2010) was used to obtain scores and locations of motifs in ISG promoters from 2000 basepairs upstream to 500 basepairs downstream of the annotated transcription start site. Promoter region diagrams were created in Inkscape to showing the relative location of sequences matching the indicated motif. KEGG and Reactome pathway enrichment values were obtained from gene set analysis using the findMotifs.pl function and selected based on differential enrichment between ISG modules.

### TR-FRET phospho-STAT1 assay

Organoids were cultured as described and treated with IFN at the indicated concentrations. Phospho-STAT1 (Y701) was detected using the LANCE *Ultra* TR-FRET kit (Perkin Elmer, #TRF4028C500). After the indicated IFN exposure times, organoid culture wells were rinsed with cold PBS-EDTA, aspirated, and 60 µL 1.25X + 25 µL 1X TR-FRET lysis buffer was added. Plates were rocked vigorously at room temperature for 30 minutes, and lysate was frozen and stored at −80 °C. Samples were then thawed, assayed in 384-well plates (Greiner, flat white) following kit instructions with a 96 hour incubation. Plates were read on a Tecan SPARK plate reader using monochromator excitation (320 nm, 10 nm bandwidth) and reading emission at 665 nm (665 nm, 8 nm bandwidth) and 615 nm (filter 620 nm, 10 nm bandwidth) with a 100 μs lag and 400 μs signal integration, set to optimal gain with a Z-distance of 20,000 μm. Data are presented as ratio of fluorescence intensity at 665/615 nm.

### Flow cytometry

Organoids were seeded at 50-100,000 cells per well and cultured for two days. For apoptosis induction experiments, organoids were treated for four hours with 1,000 U/mL of indicated IFN types, followed by treatment for 20 hours with 100 ng/mL TNFα. Organoids were dissociated with trypsin-EDTA (Gibco # 25200114) for 20-25 minutes with vigorous trituration every 5 min. Cells were then resuspended in 10 µM CellEvent Caspase 3/7 activity reporter in PBS (Invitrogen, #C10723) and incubated for 30 minutes at 37 °C, stained with ZombieAqua live/dead (BioLegend, #423102; 1:100) in PBS for 20 minutes on ice, and fixed in 2% paraformaldehyde for 15 minutes. For phospho-STAT1 experiments, organoids were treated for one hour with indicated type and concentration of IFNs, dissociated and stained with ZombieAqua live/dead as above, fixed in paraformaldehyde (BioLegend Cat 420801), permeabilized with ice-cold True-Phos™ Perm Buffer (BioLegend Cat 425401), and stained with anti-pSTAT1-PE (1:5 dilution, clone 4a, BD cat#612564). All flow cytometry was performed on a Beckman Coulter Cytoflex instrument.

### Microscopy

Organoids were cultured as described at 20,000 cells per well with apoptosis induction as described for flow cytometry. Organoids were recovered from Matrigel with cell recovery solution (Corning) at 4 °C with rocking for 30-45 min. Cells were stained as previously described (Pott et al., 2018; Van Winkle et al., 2020). Briefly, organoids were fixed with 3.7% paraformaldehyde for 10 min. at room temperature, permeabilized with ice-cold methanol at −20 °C for 20 min., blocked for 60 min. at room temperature (PBS with 5% normal goat serum, 5% bovine serum albumin, and 0.5% saponin), stained with anti-cleaved caspase 3 (Cell Signaling Technology #9661S; 1:400), anti-E-cadherin (Becton Dickinson #610182; 1:400) in staining solution (PBS with 1% normal goat serum, 1% bovine serum albumin, and 0.5% saponin) at room temperature for 4 hours, stained with anti-cleaved caspase 3 secondary antibody (goat anti-rabbit Alexa Fluor 647, Thermo Fisher; 1:400) and anti-Ecadherin secondary antibody (goat anti-mouse Alexa Fluor 555, Thermo Fisher; 1:400) for one hour at room temperature. Organoids were counterstained with DAPI (300 nM) for 5 minutes, and placed on slides with Prolong Gold antifade (Thermo Fisher). Micrographs were obtained under 20X magnification on a Zeiss ApoTome2 system on an Axio Imager, with a Zeiss AxioCam 506 camera. The CC3-positive area was measured and normalized to E-cadherin positive area (total organoid area) using ImageJ.

### RNA sequencing

The quality of the RNA samples was assessed using the TapeStation system (Agilent), and mRNA sequencing libraries were prepared using a stranded polyA+ library prep kit (Illumina). Barcoded samples from IEC organoids were separately prepared and pooled. Single-read sequencing was performed using the Illumina NovaSeq platform through the Massively Parallel Sequencing Shared Resource at OHSU.

### Sequencing analyses

Adaptor-trimmed reads were mapped to the human genome (GRCh38.89) using the STAR aligner (Dobin et al., 2013), and mapping quality was evaluated using RSeQC (Wang et al., 2012) and MultiQC (Ewels et al., 2016). One sample (Subject 005, Sigmoid colon, IFN-III-stimulated) was excluded by QC with majority of reads too short for mapping. All remaining samples had between 62 million and 85 million uniquely mapped reads with similar distributions across genomic features and uniform gene body coverage. Read counts per gene were determined using the featureCounts program (Liao et al., 2014), and differential expression analysis was performed using DEseq2 (Love et al., 2014), with a multi-factor design to control for effects of subject and biopsy site (∼ Subject + Site + Treatment + Subject:Treatment). PCA was performed on DEseq2 regularized logarithm (rlog)-transformed data. Heat maps were generated using rlog-transformed data normalized to the mean of untreated control samples; heat map clustering is based on Euclidean distance.

## Data availability

RNA sequencing data obtained in this study have been deposited in the NCBI Gene Expression Omnibus under GEO series accession number GSEXXXXX

## Contributions

DAC: conceptualization, methodology, investigation, analysis, funding acquisition, writing and editing, supervision, project administration, visualization, data curation. JVW: investigation, analysis, visualization, editing. EV: investigation, visualization. SD: funding acquisition. ER: resources. MS: resources. VT: resources. TJN: conceptualization, methodology, investigation, analysis, funding acquisition, editing, supervision, visualization.

## Ethics Statement

This study was approved by the Institutional Review Board of OHSU, protocol #00020848. Written informed consent was obtained from all participants recruited in this study.

## Acknowledgements

We thank the following OHSU core facilities for technical support, sample management, and study coordination: the Oregon Clinical and Translational Research Institute (OCTRI), the Integrated Genomics Laboratory, the Flow Cytometry Core, the Advanced Light Microscopy Core, and the OHSU Knight Cancer BioLibrary. This project was supported by the National Center for Advancing Translational Sciences (NCATS), National Institutes of Health, through Grant Award Number UL1TR002369 to OCTRI. D.A.C. was supported by NIH grant T32-AI007472, the N.L. Tartar Trust (OHSU), and the Medical Research Foundation of Oregon (OHSU). T.J.N. was supported by NIH grant R01-AI130055 and by the OHSU School of Medicine Faculty Innovation Fund. S.D. was supported by the N.L. Tartar Trust (OHSU). The funders had no role in study design, data collection and interpretation, or the decision to submit the work for publication.

